# Reorganization of Structural Connectivity in the Brain Supports Preservation of Cognitive Ability in Healthy Aging

**DOI:** 10.1101/2023.10.25.564045

**Authors:** Josh Neudorf, Kelly Shen, Anthony R. McIntosh

**Affiliations:** Institute for Neuroscience and Neurotechnology, Simon Fraser University, Burnaby, Canada; Department of Biomedical Physiology and Kinesiology, Faculty of Science, Simon Fraser University, Burnaby, Canada

**Keywords:** structural connectivity, diffusion-weighted magnetic resonance imaging, graph theory, nodal efficiency, local efficiency, healthy aging, fluid intelligence

## Abstract

The global population is aging rapidly, and a research question of critical importance is why some older adults suffer tremendous cognitive decline while others are mostly spared. Past aging research has shown that older adults with spared cognitive ability have better local short-range information processing while global long-range processing is less efficient. We took this research a step further to investigate whether the underlying structural connections, measured in vivo using diffusion magnetic resonance imaging (dMRI), show a similar shift to support cognitive ability. We analyzed the structural connectivity streamline probability (representing the probability of connection between regions) and nodal efficiency and local efficiency regional graph theory metrics to determine if age and cognitive ability are related to structural network differences. We found that the relationship between structural connectivity and cognitive ability with age was nuanced, with some differences with age that were associated with poorer cognitive outcomes, but other reorganizations that were associated with spared cognitive ability. These positive changes included strengthened local intrahemispheric connectivity and increased nodal efficiency of the ventral occipital-temporal stream, nucleus accumbens, and hippocampus for older adults, and widespread local efficiency primarily for middle-aged individuals.

**Author Summary:** We utilized network neuroscience methods to investigate why some older adults suffer tremendous cognitive decline while others are mostly spared. Past functional research found that older adults with spared cognitive ability have better local short-range information processing while global long-range processing is less efficient. We took this research a step further to investigate whether structural connectivity reorganizes to preserve cognitive ability. We analyzed age and fluid intelligence as a function of structural connectivity and regional graph theory measures using partial least squares. Some differences with age were associated with poorer cognitive outcomes, but other reorganizations spared cognitive ability. Beneficial reorganizations included strengthened local intrahemispheric connectivity and increased nodal efficiency of focal regions for older adults, as well as widespread increased local efficiency for middle-aged individuals.

---

The global population is aging rapidly (Beard et al., 2016). We are at a crucial juncture for understanding and addressing cognitive decline in this population, as emphasized by the United Nations General Assembly’s declaration that the decade from 2021 to 2030 is the Decade of Healthy Aging, calling for action to improve the health and well-being of the aging population (United Nations, 2022). Cognitive decline during aging is not an isolated concern, as it has also been shown to predict health outcomes years later (Nelson et al., 2020). This cognitive decline has been related to structural atrophy in the brain including cortical thinning (Salat et al., 2004) and myelin degradation (Bartzokis, 2004). However, research has also identified functional changes in how older adults process information that has been explained as a functional adaptation to the negative structural changes occurring (Cabeza et al., 2002; Davis et al., 2008).

Although changes in brain structural and functional connectivity have been associated with brain aging (Coelho et al., 2021; Damoiseaux, 2017; Pur et al., 2022), a demonstration that aspects of the structural connectivity network may *support* cognitive ability in older age represents an important gap in the current literature. There is, in many cases, an implicit assumption that changes with age impact an individual negatively, but by including cognitive measures in addition to age in our models we can investigate whether there are connectivity changes that impact cognitive ability positively. This idea has been supported by electroencephalography (EEG) functional research showing that local complexity (involved in short-range, intrahemispheric interactions) increases with older age, while global complexity (involved in longer range, interhemispheric interactions) decreases. Interestingly, this pattern was associated with better cognitive ability in older adults (Heisz et al., 2015). This supports the notion that not all brain changes in older age are negative, but rather that there may be reorganizations that help the brain to retain cognitive ability in the face of other physiological brain changes.

Functional connectivity represents a valuable measure of how regions work in tandem in the brain, while structural connectivity represents the physical, observable connections that evolve over the lifespan. This structural connectivity network in the brain ultimately constrains the functional states that the brain can occupy (Gu et al., 2015), and continuing research has led to growing estimates of the extent to which functional activity and functional connectivity can be accounted for based on the underlying structural connectivity (Ekstrand et al., 2020; Goñi et al., 2014; Hagmann et al., 2008; Honey et al., 2010; Neudorf et al., 2020, 2022; Rosenthal et al., 2018; Sarwar et al., 2021; Schirner et al., 2018). Knowing where there are physical changes in the brain network with age could allow for valuable interventions and prevention strategies for older adults. The whole-brain network effects of these changes are vast, varied, and challenging to assess, but graph theory approaches (Sporns, 2018) have allowed the network neuroscience research community to make great strides towards understanding the brain when viewed as a network.

We will investigate the relationship between structural connectivity in the aging brain and cognitive measures from a number of analytic and graph theory perspectives. The link between connectivity and cognitive ability in older age has been investigated using whole-brain graph theory measures of structural and functional connectivity, but structural connectivity was not associated with changes in fluid intelligence (Madden et al., 2020). In many cases, research has focused on single whole-brain measures of connectivity rather than studying the heterogeneous changes that may occur in different regions. We address the limitations of the single whole-brain measure approach by investigating the connection-specific changes and regional graph theory measures to understand the characteristics throughout the network that hinder or support cognitive ability in older age. We will investigate how structural connectivity streamline probability, representing the probability of connection between regions, are associated with cognitive ability and age, before investigating this relationship using regional graph theory measures of nodal and local efficiency. We hypothesize that this relationship will be nuanced as it is for functional analyses (Heisz et al., 2015), including some changes that are detrimental to cognitive ability, but also changes that help to spare cognitive ability in older age.

## Methods

Data came from the Cambridge Centre for Ageing and Neuroscience (Cam-CAN; Shafto et al., 2014) dataset. This data was collected in compliance with the Helsinki Declaration, and was approved by the local ethics committee, Cambridgeshire 2 Research Ethics Committee (reference: 10/H0308/50). The T1-weighted MPRAGE sequence was performed using a repetition time (TR) of 2250 ms and echo time (TE) of 2.99 ms, with a flip angle of 9°, field of view (FOV) of 256×240×192 mm, and 1×1×1 mm voxel size. The diffusion-weighted imaging was performed using a twice-refocused sequence with a TR of 9100 ms, TE of 104 ms, FOV of 192×192 mm, and voxel size of 2×2×2 mm, with 66 axial slices having a b-value of 1000, 66 axial slices having a b-value of 2000, and 66 axial slices having a b-value of 0. The structural connectivity (SC) measures of streamline probability and distance were calculated from the diffusion-weighted magnetic resonance imaging (dMRI) data using the TVB-UK Biobank pipeline (Frazier-Logue et al., 2022), which uses probabilistic tractography (*FSL bedpostx* to fit the probabilistic model and *probtrackx* to perform tractography; Hernandez-Fernandez et al., 2019; Jenkinson et al., 2012). The SC streamline probability is the number of connecting streamlines identified by the tractography divided by the total number of possible connections (i.e., normalized by the size of the region) and represents the probability of connection between all combinations of the 218 regions of interest in a combined atlas of the Schaefer 200 region atlas (Schaefer et al., 2018) and the subcortical Tian atlas (Tian et al., 2020). The SC distance was calculated as the mean distance of the streamlines between the regions of interest, taking into account the curvature of those streamlines (as calculated by *probtrackx*). The subcortical regions were comprised of regions from the Tian Scale 1 atlas excluding the hippocampus. For the hippocampus, the Scale 3 atlas was used with the two head divisions collapsed into a single parcel. The globus pallidus was excluded due to a large number of subjects without any detectible connections to or from this region, resulting in a total of 18 subcortical regions. The SC matrices were consistency thresholded (at least 50% of participants have the connection) and participants’ data were excluded if they did not have behavioral data, had regions with no connections, or had SC density (number of non-zero connections divided by the total number of possible connections) 3 standard deviations (SD) or more away from the mean (retained N = 594 from the total of 656 participants in the dataset). The variables of interest, age and fluid intelligence, were not significantly associated with the total SC streamline probability when analyzed in a multilinear model (model *R^2^* = .006, *p* = .193). Total SC streamline probability was significantly different between male and female participants, *t*(592) = -3.064, *p* = .002. We included sex as a variable in subsequent analyses and found that this difference did not alter the effects of interest.

The Cattell Culture Fair fluid intelligence score (Cattell & Cattell, 1973) was used to measure cognitive ability. This measure of cognitive ability declines significantly with age in this population, *R*(592) = -.651, *p* < .001 (see Figure 1), so factors that counteract this decline represent a sparing of cognitive ability.

**Figure 1.**
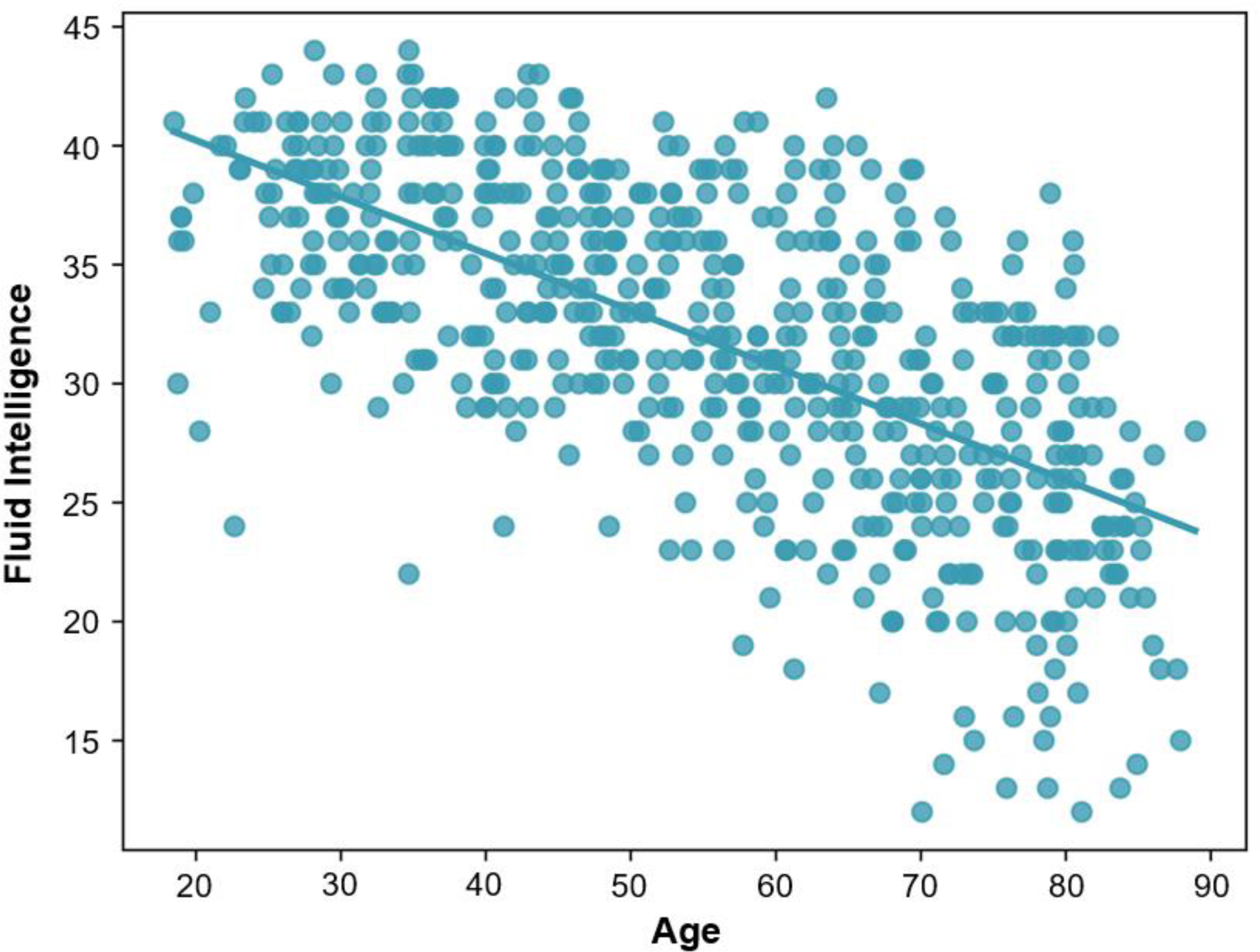
Relationship between age and fluid intelligence as measured by the Cattell Culture Fair fluid intelligence score. Fluid intelligence decreases significantly with age, *R*(592) = -.651, *p* < .001.

Participant ages ranged from 18.50 to 88.92 (mean = 55.414, *SD* = 18.090). Younger adults (YA; age < 50) ranged from 18.50 to 49.92 years (mean = 36.966, *SD* = 8.385, *N* = 244, 131 female, 113 male) and older adults (OA; age > 50) ranged from 50.17 to 88.92 years (mean = 68.275, *SD* = 10.163, *N* = 350, 170 female, 180 male).

Multivariate partial least squares (PLS) analysis (McIntosh & Lobaugh, 2004) was used to identify latent variables (LVs), each containing weights that describe the relationship of all connections or regional graph theory measures with age and fluid intelligence. Models including sex as a group variable were examined, but produced no significant group effect so they were not included in the reported analyses. PLS uses singular value decomposition to project the data matrix onto orthogonal LVs (similar to principal component analysis; PCA). The significance of each identified LV is determined via permutation testing, whereby the order of the data is randomized over a large number of iterations so that the independent and dependent variables are no longer paired to one another, in order to obtain a permutation p-value representing the significance of that LV. We report only the most reliable PLS weights as determined by bootstrap resampling, which is used to calculate bootstrap ratios (BSR). The bootstrap ratio indicates the reliability of the PLS weights and is calculated as the ratio of the PLS weights (saliences) and their standard errors as determined by bootstrap resampling (Kovacevic et al., 2013; McIntosh & Lobaugh, 2004). The BSRs are analogous to a Z-score, and as such a value exceeding an absolute value of 2 is considered reliable. This procedure was performed using 1000 iterations for permutation testing and bootstrap resampling. To assess the out-of-sample reproducibility of the PLS results, a split-half approach was used selecting 2 random groups of subjects and comparing the singular values between these groups and the singular vectors between groups (as described by McIntosh, 2022). This process was repeated 1000 times to produce the Z_sval_ (for singular values) and the Z_svec_ (for singular vectors) statistics, indicating how reproducible the LV was. Null distributions generated using randomly permuted data were used to generate a null Z value (Z_sval_null_ and Z_svec_null_), which was added to the typical threshold of 2.0 in order to produce a threshold for reproducibility.

Age and fluid intelligence were examined as dependent variables, with three separate PLS analyses for SC streamline probability, as well as graph theory measures of SC-based nodal efficiency (NE) and SC-based local efficiency (LE). Nodal efficiency is calculated as the mean of the inverse shortest path length to all other regions in the network,

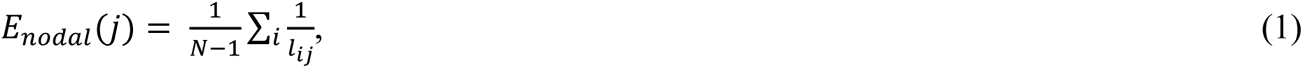

where *l*_*ij*_ is the weighted shortest path length between regions *i* and *j* (Latora & Marchiori, 2001). Nodal efficiency gives a measure of how closely connected a region is to the rest of the brain, and is high for hub regions in the brain that are important for linking regions from different modules (Achard & Bullmore, 2007; van den Heuvel & Sporns, 2013). Local efficiency is a separate measure of efficiency distinct from nodal efficiency, which is calculated as the mean of the inverse shortest path length for the subnetwork of connected regions after first removing the region of interest,

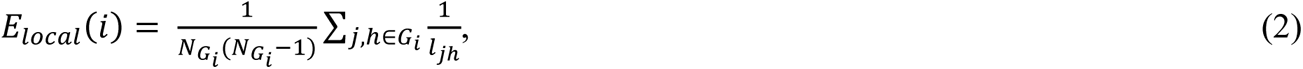

where *G*_*i*_ is the subgraph/subnetwork made up of the neighboring regions connected to region *i*, *N*_*Gi*_ is the number of regions in this subgraph, and *l*_*j*ℎ_is the weighted shortest path length between regions *j* and ℎ in this subgraph (Latora & Marchiori, 2001). Local efficiency is thus a measure of how well connected the neighbors of a region are, and can been conceptualized as a measure of fault tolerance, or robustness against disruption to the network (Fornito et al., 2016).

Once important regions were identified based on these graph theory measures, a supplementary rolling correlation analyses was conducted on age bins of 10 consecutive years to identify critical periods during which these graph theory measures were most highly related to cognitive ability. This type of analysis is implemented in many statistical packages, and has been used in the analysis of financial time-series data (e.g., Ilalan & Pirgaip, 2019). The rolling correlation analysis consisted of selecting bins of participants with a range of 10 years of age and calculating the Pearson’s correlation coefficient (*R*) between the mean graph theory measure and the fluid intelligence score. Bins of this size resulted in a mean number of participants in each bin of 91.532, with a standard deviation of 14.330, ranging from 42 to 111. Of the 62 total bins, 54 (87%) had over 80 participants, and bins with fewer participants were largely situated in the 7 youngest age bins. The confidence intervals of these correlation coefficients was calculated using the bias-corrected and accelerated bootstrap method (Efron, 1987), in order to determine whether the confidence intervals did not show extreme fluctuations across age bins.

A number of analysis decisions were made in response to our findings in order to better understand these relationships. The PLS analysis of SC with all subjects included revealed a LV associated with connections that supported cognitive ability in YA, as well as a LV that supported cognitive ability in OA. However, the PLS analysis of nodal efficiency and local efficiency identified only a single LV, so separate analyses were conducted on the YA and OA groups to better understand the differences in the relationship with fluid intelligence between these age groups. These YA and OA nodal efficiency analyses revealed interesting differences in the pattern of regions related to fluid intelligence, whereby OA had noticeably more regions supporting fluid intelligence than YA. This motivated the additional rolling correlation analysis focusing on the regions that were identified as important in OA to more quantitatively test the hypothesis that these patterns indicate a growing importance of nodal efficiency into older adulthood. In the same vein, the YA and OA local efficiency analyses motivated a further rolling correlation analysis as OA related regions were exclusively negatively related to fluid intelligence, whereas the YA related regions were primarily positively related to fluid intelligence. As in the nodal efficiency analysis, we focused on the regions that were positively associated with fluid intelligence, which was found in the YA group, and examined this relationship across the lifespan with the rolling correlation analysis.

Finally, as mentioned, the OA local efficiency analysis revealed that local efficiency was exclusively negatively related to fluid intelligence, and that this pattern implicated widespread regions across the whole brain. We investigated this finding further by examining the mean local efficiency of these regions in a multilinear model with whole-brain independent variables of small-worldness, SC total, and SC distance (as well as age and sex) to determine how this widespread pattern of local efficiency may be related to differences in the network as a whole. A small-world network has high clustering coefficient and low shortest path length as opposed to a random network (low shortest path length but low clustering coefficient). There are many examples of real-world networks that have these small-world properties, including brain networks (Hilgetag et al., 2000; Micheloyannis, Pachou, Stam, Breakspear, et al., 2006; Micheloyannis, Pachou, Stam, Vourkas, et al., 2006; Salvador et al., 2005; Stam et al., 2007). The clustering coefficient measures how likely two neighbors of a region are to also be connected to one another, and supports the ability for subnetworks of the brain to integrate information and be specialized (Onnela et al., 2005). On the other hand, a low shortest path length (Dijkstra, 1959) indicates that there are short routes between regions in the brain in general, which allows the specialized subnetworks to communicate across the whole brain efficiently (Fornito et al., 2016). Research has identified multiple ways in which small-worldness supports organized functional activity, by improving the capacity for functional segregation as well as integration (Sporns et al., 2000; Sporns & Zwi, 2004), synchronization and propagation of information (Barahona & Pecora, 2002; Hong et al., 2002), computational power (Lago-Fernández et al., 2000), and more (see reviews by Bassett & Bullmore, 2006; Fornito et al., 2016).

## Data and Code Availability Statement

The data used in this analysis are available at https://cam-can.org. Code used to produce these analyses are available at https://github.com/neudorf/sc-efficiency-aging.

## Results

### SC Streamline Probability PLS

The PLS analysis of SC streamline probability identified 2 significant LVs. LV1 (permutation *p* < .001; Z_sval_ = 34.736 > 2 + Z_sval_null_ = 2.046; Z_svec_ = 95.273 > 2 + Z_svec_null_ = 3.231) was negatively correlated with age, *R* = -.813, *95% CI* = [-.852, -.811], and positively correlated with fluid intelligence, *R* = .615, *95% CI* = [.613, .682], indicating that more streamlines in positively weighted connections related to better cognitive ability in younger age. On the other hand, negative connection weights indicate that fewer streamlines in these connections related to better fluid intelligence in younger age. LV2 (permutation *p* = .045; Z_sval_ = 2.107 > 2 + Z_sval_null_ = 2.012; Z_svec_ = 2.601 < 2 + Z_svec_null_ = 3.294) was positively correlated with both age, *R* = .313, *95% CI* = [.067, .371] and fluid intelligence, *R* = .300, *95% CI* = [.283, .512], indicating that more streamlines in connections with positive weights related to better fluid intelligence in older age. Negative weights, meanwhile, indicate that fewer streamlines in these connections related to better fluid intelligence in older age.

### Latent Variable 1: Positive Weights

The largest positive weights from LV1 identified a subnetwork with roughly equal numbers of intrahemispheric (41.7%; 13.9% LH and 27.8% RH) and interhemispheric (58.3%) connections. Notable regions with a degree (number of connections) of at least 3 included bilateral medial parietal cortex, bilateral postcentral gyrus, LH dorsal prefrontal cortex (dPFC), LH precuneus, RH medial prefrontal cortex (mPFC), and RH hippocampus tail (see Figure 2). Of these regions, the two superior frontal regions, hippocampus, and precuneus have been previously identified as rich-club regions in the brain (highly connected hub regions that are also highly connected to one another; van den Heuvel & Sporns, 2011).

**Figure 2.**
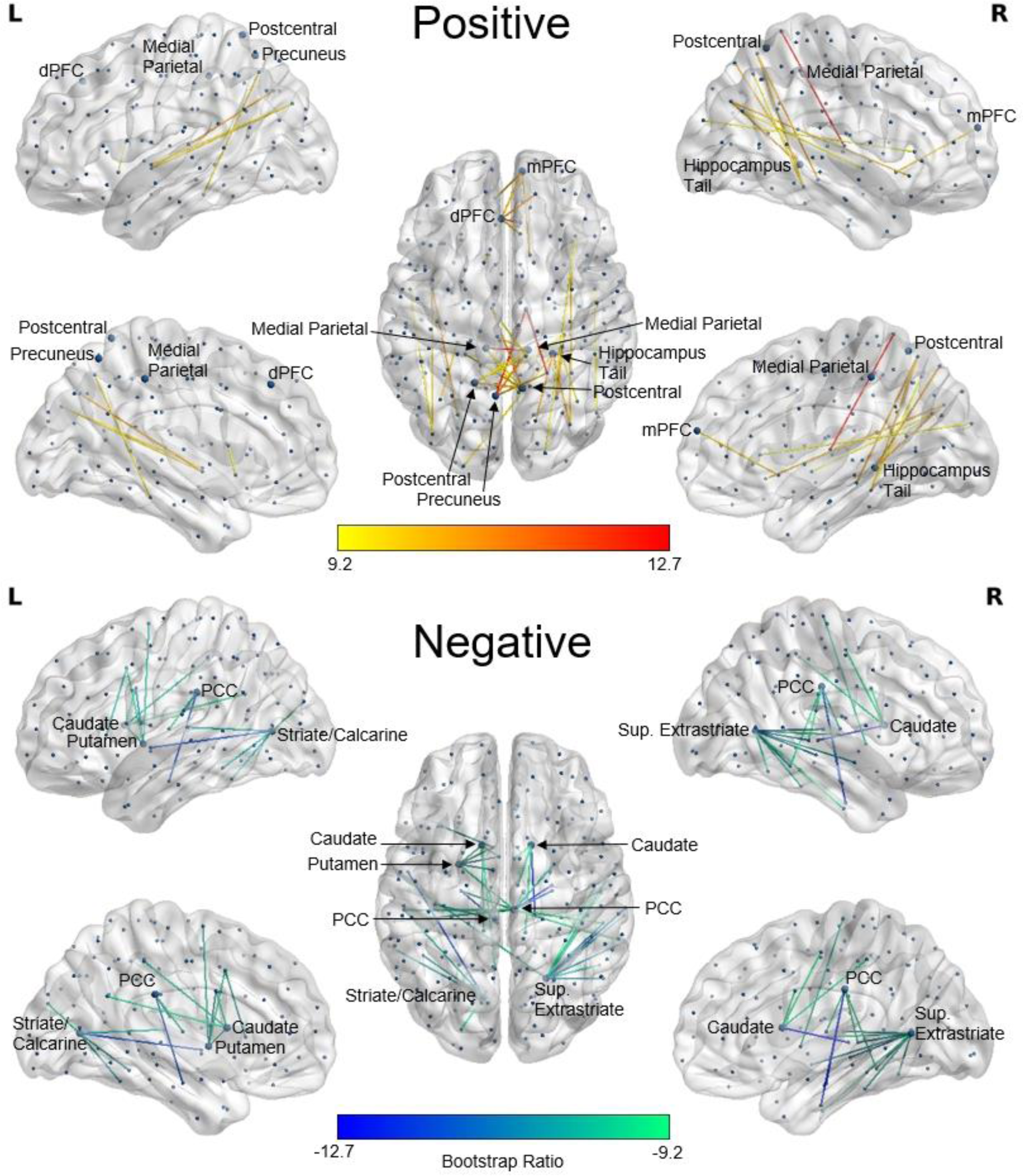
SC PLS analysis of age and fluid intelligence. Shown are LV1’s most reliable connections as identified by bootstrap resampling. Connection color corresponds to the value of the BSR for that connection. Positive BSRs (top) indicate where stronger connections are related to better fluid intelligence in younger age, whereas negative BSRs (bottom) indicate where fewer connections are related to better fluid intelligence in younger age. The BSR threshold of ±9.2 was selected to keep the number of displayed connections comparable for LV1 and LV2. Highly connected regions with degree (number of connections) greater than 3 are labeled. Figure produced in BrainNet Viewer (Xia et al., 2013).

### Latent Variable 1: Negative Weights

The largest negative weights from LV1 identified a subnetwork made up of mostly intrahemispheric connections (87.8%; 39.0% LH and 48.8% RH), with the exception of 5 (12.2%) interhemispheric connections (1 to the RH superior extrastriate and 4 to the RH posterior cingulate cortex; PCC). Highly implicated regions in this network include the bilateral PCC, bilateral caudate nucleus, LH striate/calcarine cortex, LH putamen, and RH superior extrastriate (see Figure 2). In combination, LH striate/calcarine and RH superior extrastriate play a role in approximately half of the connections (48.8%).

### Latent Variable 2: Positive Weights

For LV2, the largest positive weights were primarily intrahemispheric (94.1%) and split equally between the LH (47.1%) and RH (47.1%), with a small number of interhemispheric connections identified (5.9%; see Figure 3). Furthermore, these connections supporting fluid intelligence in older age were shorter in length than the connections identified by positive weights for LV1 representing connections supporting fluid intelligence earlier in the lifespan, *t*(593) = -10.062, *p* < .001, and also shorter than the connections identified by the LV2 negative weights that were negatively associated with fluid intelligence in older age, *t*(593) = -30.454, *p* < .001, based on paired samples *t*-tests.

**Figure 3.**
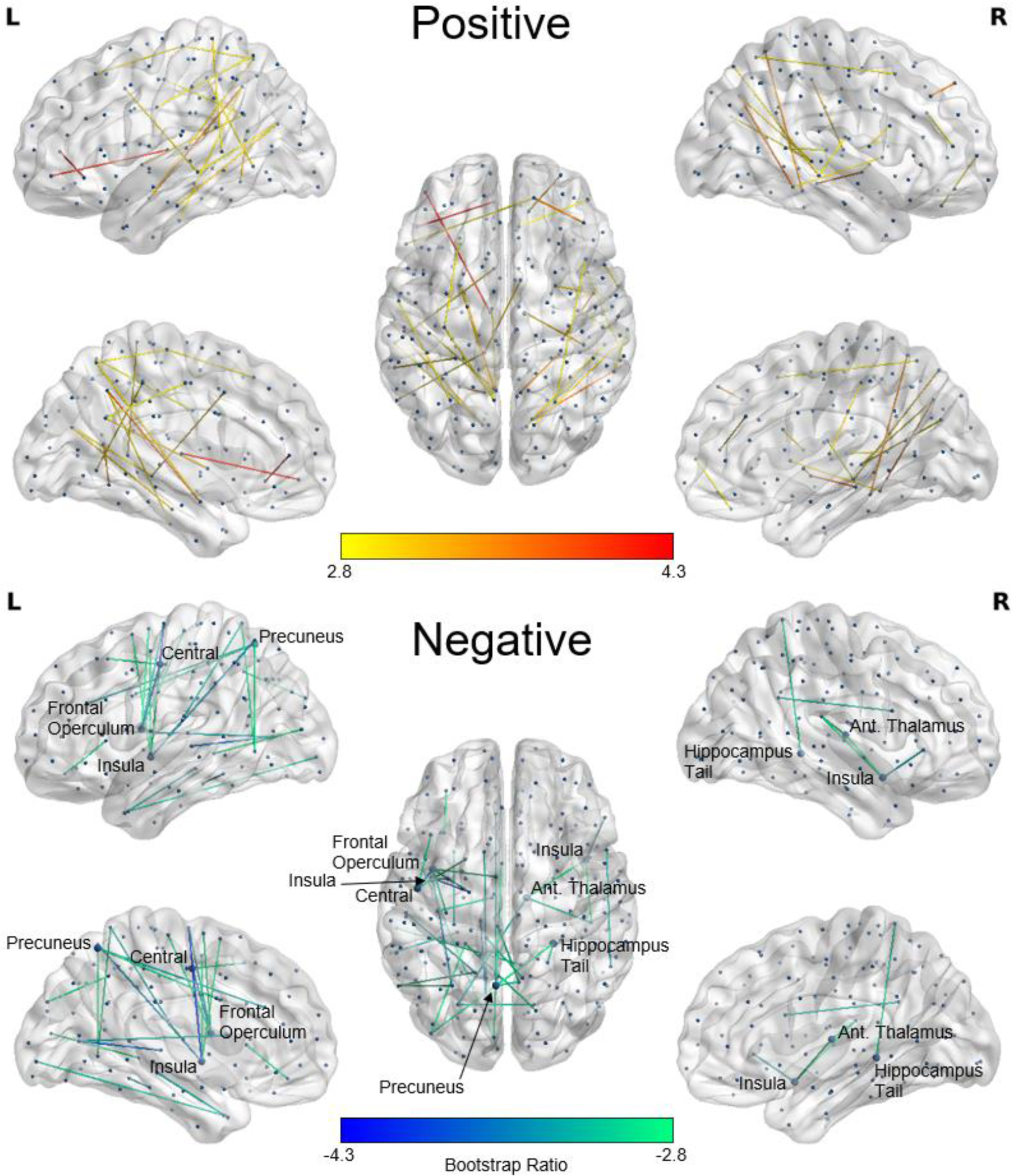
SC PLS analysis of age and fluid intelligence. Shown are LV2’s most reliable connections as identified by bootstrap resampling. Connection color corresponds to the value of the BSR for that connection. Positive BSRs (top) indicate where stronger connections are related to better fluid intelligence in older age, whereas negative BSRs (bottom) indicate where fewer connections are related to better fluid intelligence in older age. The BSR threshold of ±2.8 was selected to keep the number of displayed connections comparable for LV1 and LV2. Highly connected regions with degree (number of connections) greater than 3 are labeled.

### Latent Variable 2: Negative Weights

The largest negative weights for LV2 included mostly intrahemispheric connections (76.9%; 61.5% LH and 15.4% RH) with a smaller number of interhemispheric connections (23.1%). Highly implicated regions include the bilateral insula, LH frontal operculum, LH central sulcus, LH precuneus, RH anterior thalamus, and RH hippocampus tail (see Figure 3).

## SC Nodal Efficiency PLS

### Younger Adults

The SC nodal efficiency PLS analysis for YAs identified 1 significant LV. This LV (permutation *p* = .012; Z_sval_ = 1.769 < 2 + Z_sval_null_ = 2.036; Z_svec_ = 2.171 < 2 + Z_svec_null_ = 3.312) was negatively correlated with age, *R* = -.482, *95% CI* = [-.616, -.459], and positively correlated with fluid intelligence, *R* = .113, *95% CI* = [.047, .266], indicating that higher nodal efficiency in positively weighted regions related to better fluid intelligence in younger age. On the other hand, negative region weights indicate that lower nodal efficiency in these regions contribute to better fluid intelligence in younger age.

For these YA, PLS identified a clustered pattern of negative connection weights in the insula and lateral prefrontal cortex, indicating that less nodal efficiency is better for fluid intelligence in these regions. These regions include bilateral insula, bilateral ventrolateral and dorsolateral prefrontal cortex, and bilateral putamen. There were also positively weighted regions indicating that more nodal efficiency is better for fluid intelligence in these regions. These regions included the LH precuneus, LH striate, LH mPFC, LH hippocampus, RH striate and extrastriate, RH inferior parietal lobule (IPL), and RH temporal pole (see Figure 4; see Supplementary Figure 1 for subcortical region labels).

**Figure 4.**
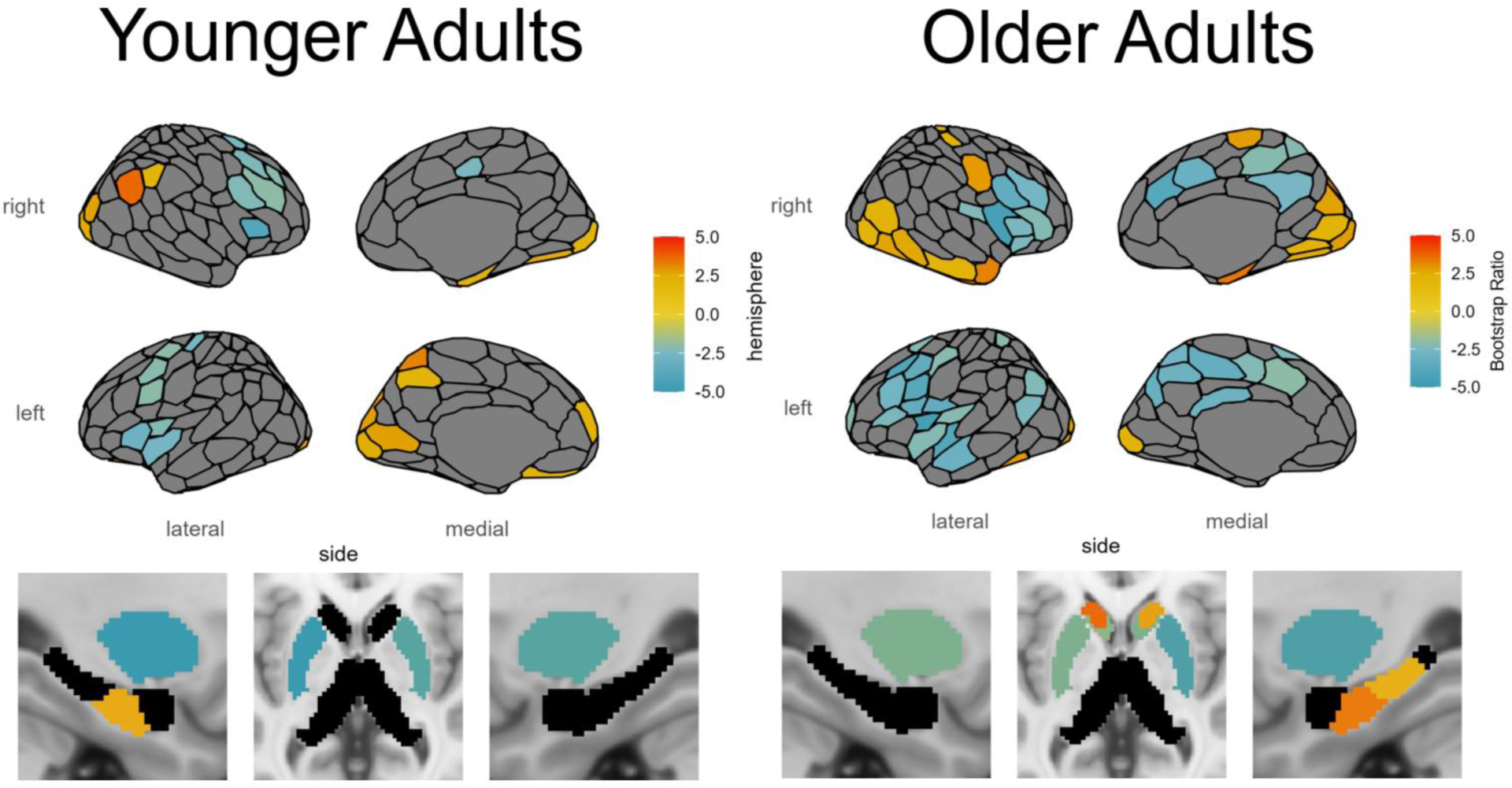
SC-based nodal efficiency PLS analysis of age and fluid intelligence. Shown are the regions identified as reliable with respect to the correlation with age and cognitive ability. Region color corresponds to the value of the BSR for that region, thresholded at ±2.0. See Supplementary Figure 1 for subcortical region labels.

### Older Adults

The PLS analysis for OAs identified 1 significant LV. This LV (permutation *p <* .001; Z_sval_ = 6.663 > 2 + Z_sval_null_ = 2.049; Z_svec_ = 6.442 > 2 + Z_svec_null_ = 3.328) was negatively correlated with age, *R* = -.467, *95% CI* = [-.632, -.428], and positively correlated with fluid intelligence, *R* = .316, *95% CI* = [.289, .482], indicating that higher nodal efficiency in positively weighted regions related to better fluid intelligence in younger age. On the other hand, negative region weights indicate that lower nodal efficiency in these regions contributed to better fluid intelligence in younger age.

For these OA, PLS identified a similar clustered pattern of negative connection weights in the insula and prefrontal cortex as was seen in the YA, but with more regions involved bilaterally in medial parietal regions, medial and lateral frontal regions, and the caudate nucleus, and greater involvement in the LH superior temporal lobe and LH auditory cortex. For the positively weighted regions indicating better fluid intelligence, PLS identified a greater number of regions than the YA, including bilateral nucleus accumbens and a striking pattern of regions in the RH, including the striate, extrastriate, and ventral occipital-temporal stream, with involvement of this ventral stream all the way to the temporal pole, as well as postcentral gyrus and hippocampus (see Figure 4).

### Rolling Correlation Analysis: Nodal Efficiency and Cognitive Ability

In order to investigate whether there may have been a critical age period during which nodal efficiency was beneficial for fluid intelligence in the regions identified as being positively related to fluid intelligence in OA, a rolling Pearson’s correlation coefficient analysis between the mean nodal efficiency in these regions and fluid intelligence was calculated with a window size of 10 years (see Figure 5). The size of confidence intervals appeared consistent across the different age bins. This analysis revealed an inverted ‘U’ shape in this relationship, and identified a critical period during which the nodal efficiency of these regions was beneficial for fluid intelligence, during the ages of approximately 55-72 years.

**Figure 5.**
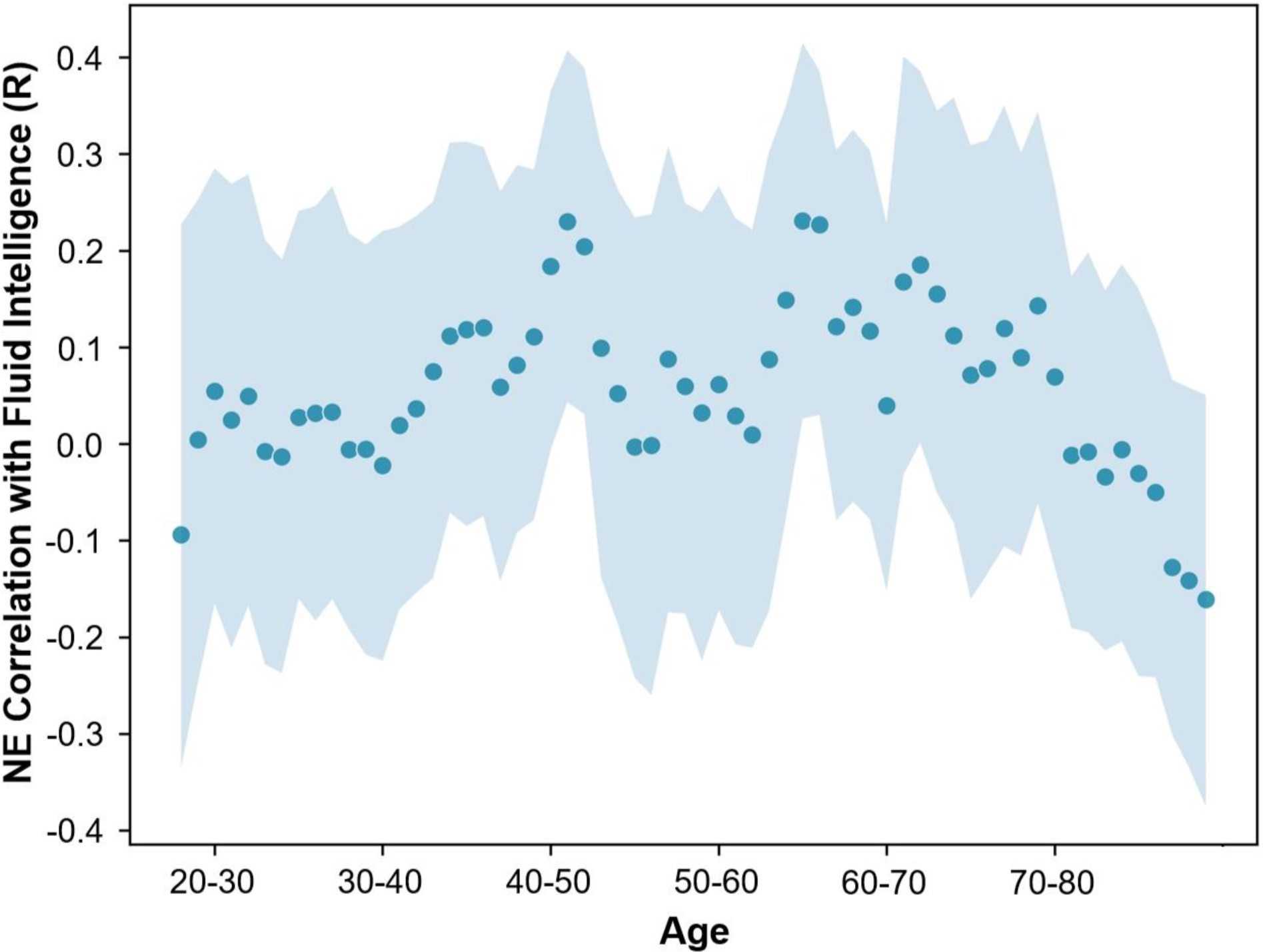
Rolling Pearson’s correlation coefficient between mean nodal efficiency and fluid intelligence for regions positively related to the PLS LV in OA for participants in age windows of 10 years. Shaded area represents the 95% CI.

### Mean Nodal Efficiency

To better understand the pattern of negatively weighted regions with respect to nodal efficiency, we also present the mean nodal efficiency values, as a representation of typical nodal efficiency in this population (see Figure 6). The pattern of higher nodal efficiency hub-like regions overlapped with those regions identified as being negatively related to fluid intelligence, especially for OA but also for YA, particularly in the insula, lateral PFC, cingulate cortex, medial parietal regions, putamen, and caudate nucleus. We can demonstrate this effect quantitatively by comparing the nodal efficiency of regions identified as being negatively related to fluid intelligence in OA to those that were positively related to fluid intelligence, *t*(593) = 79.953, *p* < .001, and to all other regions, *t*(593) = 59.736, *p* < .001, with paired samples *t*-tests. The high nodal efficiency of these regions suggests that there may be an optimal network configuration of the most hub-like regions in the brain, and that alterations to this configuration were detrimental to fluid intelligence.

**Figure 6.**
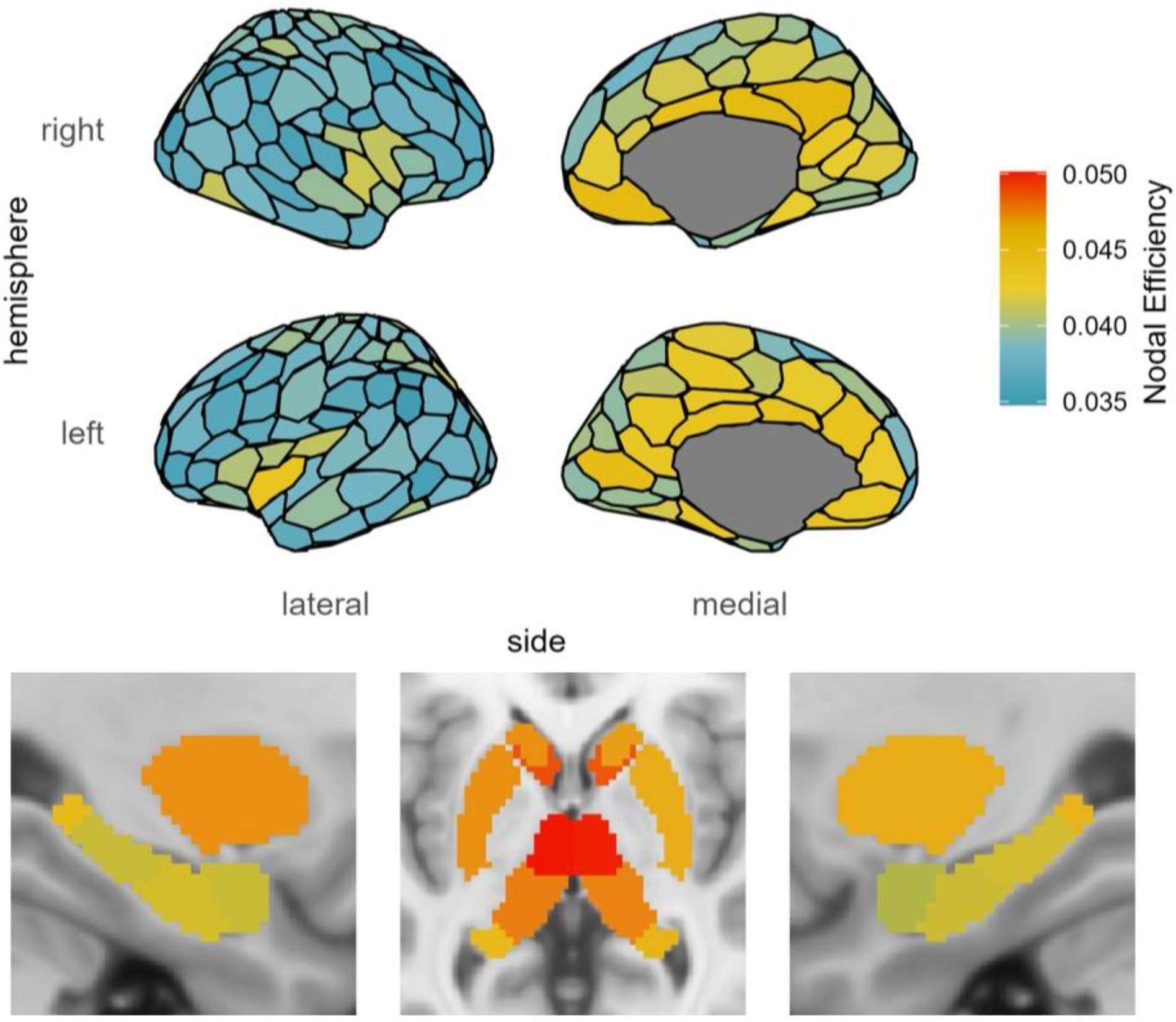
Brain map of mean nodal efficiency for all participants.

## SC Local Efficiency PLS

### Younger Adults

The SC local efficiency PLS analysis for YAs identified 1 significant LV. This LV (permutation *p* < .001; Z_sval_ = 5.112 > 2 + Z_sval_null_ = 2.025; Z_svec_ = 4.989 > 2 + Z_svec_null_ = 3.273) was negatively correlated with age (*R* = -.352, *95% CI* = [-.548, -.309]) and positively correlated with fluid intelligence (*R* = .134, *95% CI* = [.081, .263]), indicating that higher local efficiency in positively weighted regions related to better fluid intelligence in younger age. On the other hand, negative region weights indicate that lower local efficiency in these regions contributed to better fluid intelligence in younger age.

For these YA, local efficiency was positively related to fluid intelligence in the large majority of regions, including bilateral cingulate cortex, bilateral precuneus, bilateral retrosplenial cortex, bilateral medial parietal cortex, bilateral temporal and temporoparietal cortex, bilateral auditory cortex, LH hippocampus body, RH parahippocampal cortex, RH superior and inferior extrastriate, RH postcentral gyrus, RH precentral gyrus, RH dorsal prefrontal cortex, RH insula, RH posterior thalamus, and RH hippocampus, with a smaller number of negatively related regions in the bilateral orbitofrontal cortex, bilateral ventral prefrontal cortex, bilateral anterior thalamus, LH central sulcus, and LH inferior parietal sulcus (see Figure 7).

**Figure 7.**
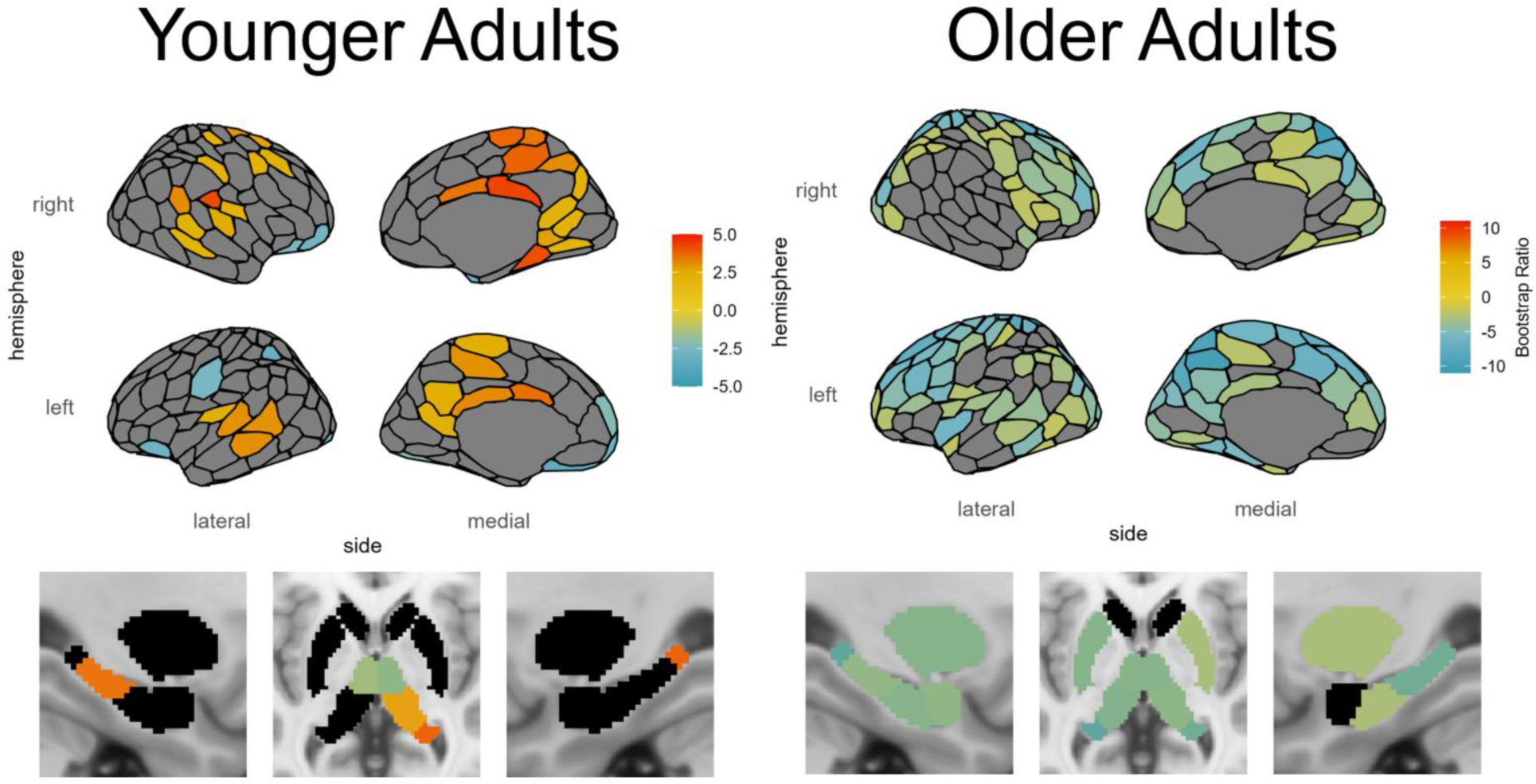
SC-based local efficiency PLS analysis of age and fluid intelligence. Shown are the regions identified as reliable with respect to the correlation with age and fluid intelligence. Region color corresponds to the value of the BSR for that region, thresholded at ± 2.0.

### Older Adults

The PLS analysis for OAs identified 1 significant LV. This LV (permutation *p <* .001; Z_sval_ = 10.033 > 2 + Z_sval_null_ = 2.003; Z_svec_ = 40.921 > 2 + Z_svec_null_ = 3.271) was negatively correlated with age, *R* = -.430, *95% CI* = [-.576, -.354], and positively correlated with fluid intelligence, *R* = .274, *95% CI* = [.230, .405], indicating that higher local efficiency in positively weighted regions related to better fluid intelligence in younger age. On the other hand, negative region weights indicate that lower local efficiency in these regions contributed to better fluid intelligence in younger age.

Unlike the YA, for these OA local efficiency was negatively related to fluid intelligence in all reliable regions, which were distributed throughout large portions of the brain (see Figure 7).

### Rolling Correlation Analysis: Local Efficiency and Fluid Intelligence

To investigate whether there may be a critical age period during which local efficiency is beneficial for fluid intelligence in the regions identified as being positively related to fluid intelligence in YA, a rolling Pearson’s correlation coefficient analysis between the mean nodal efficiency in these regions and fluid intelligence was calculated with a window size of 10 years (see Figure 8). The sizes of confidence intervals appeared consistent across the different age bins. This analysis revealed a critical period during which the nodal efficiency of these regions was beneficial for fluid intelligence, during the ages of approximately 32-51 years.

**Figure 8.**
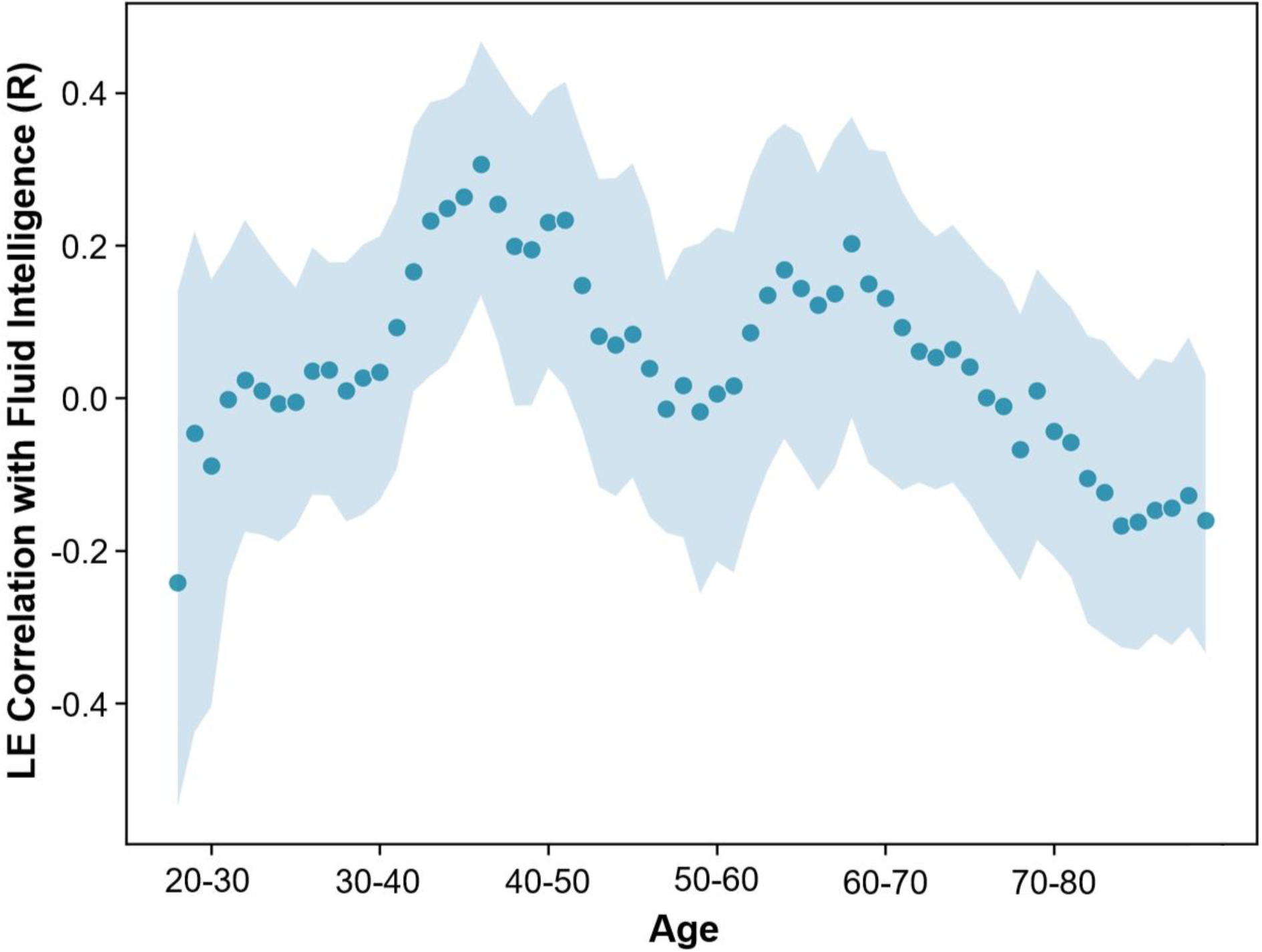
Rolling Pearson’s correlation coefficient between mean local efficiency and fluid intelligence for regions positively related to the PLS LV in YA for participants in age windows of 10 years. Shaded area represents the 95% CI.

### Whole-brain Network Effects of Local Efficiency

Mean local efficiency (in the regions identified as negatively associated with fluid intelligence in OA; see Fig. 7) had no significant relationship with age in YA, *R*(242) = -.094, *p* = .143, but a positive relationship with age in OA, *R*(348) = .461, *p* < .001 (see Figure 9).

**Figure 9.**
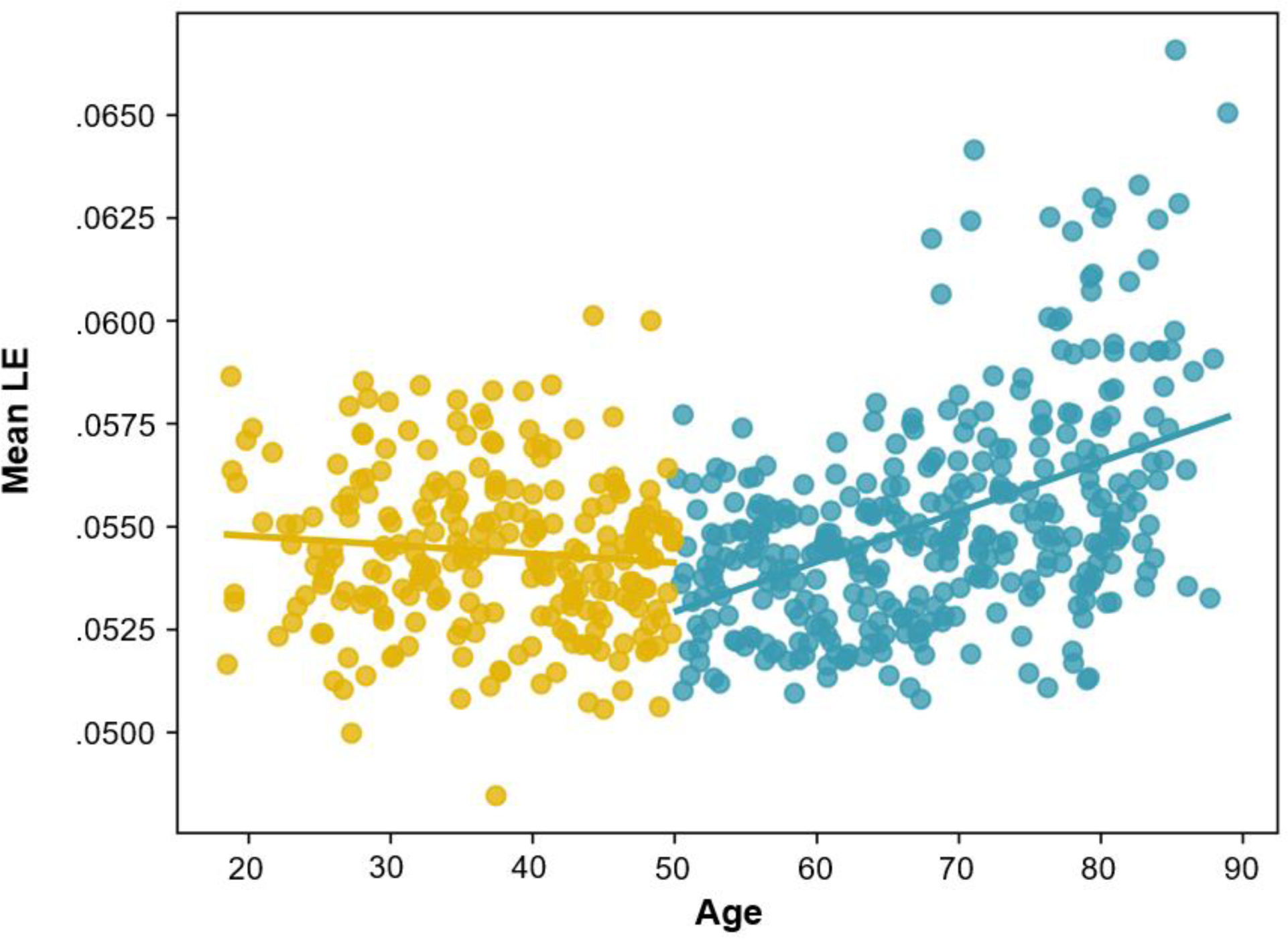
Relationship between age and mean local efficiency (in the regions identified as negatively associated with fluid intelligence in OA) for YA (in yellow; *R*(242) = -.094, *p* = .143) and OA (in blue; *R*(348) = .461, *p* < .001).

In order to understand what changes are occurring in the structural connectivity network at the whole-brain level as local efficiency increases in OA, multiple linear regression models were created with mean local efficiency (in the regions identified as negatively associated with fluid intelligence in OA) as the dependent variable, and graph theory measures of small-worldness, SC total (total of SC probabilities in the brain), and mean SC streamline distance as the independent variables.

## Model 1

Model 1 included independent variables of age, sex, small-worldness, and SC total (total of SC streamline probabilities), with the dependent variable (which will be the same for all models) of local efficiency mean for regions negatively associated with fluid intelligence in OA based on the PLS analysis (see Figure 7). For YA, local efficiency was negatively associated with age now that these other covariates have been included in the model, was associated with greater small-worldness, and was associated with greater SC total (see Table 1). For OA, local efficiency was *positively* associated with age, was associated with greater small-worldness, was associated with greater SC total, and males had lower local efficiency than females (see Table 2). These results identified commonalities as well as differences between these age groups in the way that local efficiency impacts the brain network. In both YA and OA increased local efficiency was associated with greater small-worldness. However, local efficiency was associated more strongly with small-worldness in OA (estimate value of 4.512*10^-3^ compared to 1.745*10^-3^ for YA, which was significant based on the estimates’ 95% confidence intervals; CI; see Tables 1 and 2), indicating that the relationship with small-worldness was stronger for OA than YA. Local efficiency was negatively associated with age in YA, but positively associated with age in OA, and this difference was significant based on the 95% CI of the estimate. Sex was not identified as a significant factor for YA, but OA males had lower local efficiency than females, and this difference was significant based on the 95% CI of the estimate. Furthermore, while local efficiency was positively associated with SC total for YA and OA, this relationship was stronger for OA (significant based on the 95% CI of the estimate). Finally, a much larger amount of variance was accounted for by the model in OA (*R*^2^ = .723 compared to *R*^2^ = .341 for YA), indicating that these whole-brain measures accounted for more variance in local efficiency for OA.

**Table 1.** Local Efficiency Multilinear Regression Analysis Model 1 for YA, with dependent variable local efficiency mean for regions negatively associated with fluid intelligence in OA based on the PLS analysis. Independent variables included sex, age, small-worldness, and SC total (total SC streamline probabilities). Model *R*^2^ = .341.

| Effects | Estimate [95% CI] | Std. Error | *t*-value | *p*-value |
| --- | --- | --- | --- | --- |
| Intercept | .013 [.004, .021] | .004 | 2.942 | .004 |
| Age | $-2.778 \times 10^{-5}$ [$-5.174 \times 10^{-5}$, $-3.830 \times 10^{-6}$] | $1.216 \times 10^{-5}$ | -2.285 | .023 |
| Sex (Male) | $1.344 \times 10^{-4}$ [$-2.657 \times 10^{-4}$, $5.346 \times 10^{-4}$] | $2.031 \times 10^{-4}$ | .662 | .509 |
| Small-worldness | $1.745 \times 10^{-3}$ [$1.342 \times 10^{-3}$, $2.147 \times 10^{-3}$] | $2.043 \times 10^{-4}$ | 8.539 | <.001 |
| SC Total | $1.451 \times 10^{-4}$ [$1.126 \times 10^{-4}$, $1.777 \times 10^{-4}$] | $1.652 \times 10^{-5}$ | 8.785 | <.001 |

**Table 2.** Local Efficiency Multilinear Regression Analysis Model 1 for OA, with dependent variable local efficiency mean for regions negatively associated with fluid intelligence based on the PLS analysis. Independent variables included sex, age, small-worldness, and SC total (total SC streamline probabilities). Model *R*^2^ = .723.

| Effects | Estimate [95% CI] | Std. Error | *t*-value | *p*-value |
| --- | --- | --- | --- | --- |
| Intercept | -.019 [-.026, -.012] | .004 | -5.516 | <.001 |
| Age | $6.857 \times 10^{-5}$ [$5.327 \times 10^{-5}$, $8.388 \times 10^{-5}$] | $7.782 \times 10^{-6}$ | 8.812 | <.001 |
| Sex (Male) | $-6.023 \times 10^{-4}$ [$-9.145 \times 10^{-4}$, $-2.901 \times 10^{-4}$] | $1.587 \times 10^{-4}$ | -3.795 | <.001 |
| Small-worldness | $4.512 \times 10^{-3}$ [$4.141 \times 10^{-3}$, $4.883 \times 10^{-3}$] | $1.886 \times 10^{-4}$ | 23.924 | <.001 |
| SC Total | $2.100 \times 10^{-4}$ [$1.844 \times 10^{-4}$, $2.356 \times 10^{-4}$] | $1.300 \times 10^{-5}$ | 16.154 | <.001 |

## Model 2

Model 2 included independent variables of age, sex, small-worldness, and mean SC distance (mean length of the structural connections in the brain, taking into account the curvature of the connections), differing from Model 1 by the substitution of SC distance for SC total. The results mirrored those found in Model 1, but additionally we found that SC distance was negatively associated with local efficiency for both YA (see Table 3) and OA (see Table 4), and this effect was stronger for OA than YA (significant based on the 95% CI of the estimate). We also found that YA males had higher local efficiency than females (whereas this was not significant in Model 1).

**Table 3.** Local Efficiency Multilinear Regression Analysis Model 2 for YA, with dependent variable local efficiency mean for regions negatively associated with fluid intelligence based on the PLS analysis. Independent variables included sex, age, small-worldness, and SC distance (mean of all SC streamline distances). Model *R*^2^ = .171.

| Effects | Estimate [95% CI] | Std. Error | *t*-value | *p*-value |
| --- | --- | --- | --- | --- |
| Intercept | .060 [.054, .066] | .003 | 19.337 | <.001 |
| Age | -2.431*10<sup>-5</sup> [-5.135*10<sup>-5</sup>, 2.727*10<sup>-6</sup>] | 1.372*10<sup>-5</sup> | -1.771 | .078 |
| Sex (Male) | 5.495*10<sup>-4</sup> [3.791*10<sup>-5</sup>, 1.061*10<sup>-3</sup>] | 2.597*10<sup>-4</sup> | 2.116 | .035 |
| Small-worldness | 9.837*10<sup>-4</sup> [5.152*10<sup>-4</sup>, 1.452*10<sup>-3</sup>] | 2.378*10<sup>-4</sup> | 4.136 | <.001 |
| SC Distance | -1.380*10<sup>-4</sup> [-2.155*10<sup>-4</sup>, -6.064*10<sup>-5</sup>] | 3.929*10<sup>-5</sup> | -3.513 | <.001 |

**Table 4.** Local Efficiency Multilinear Regression Analysis Model 2 for OA, with dependent variable local efficiency mean for regions negatively associated with fluid intelligence based on the PLS analysis. Independent variables included sex, age, small-worldness, and SC distance (mean of all SC streamline distances). Model *R*^2^ = .573.

| Effects | Estimate [95%CI] | Std. Error | *t*-value | *p*-value |
| --- | --- | --- | --- | --- |
| Intercept | .053 [.048, .058] | .003 | 20.171 | <.001 |
| Age | 7.295*10<sup>-5</sup> [5.385*10<sup>-5</sup>, 9.204*10<sup>-5</sup>] | 9.709*10<sup>-6</sup> | 7.513 | <.001 |
| Sex (Male) | -3.059*10<sup>-5</sup> [-4.803*10<sup>-4</sup>, 4.191*10<sup>-4</sup>] | 2.286*10<sup>-4</sup> | -.134 | .894 |
| Small-worldness | 2.458*10<sup>-3</sup> [1.956*10<sup>-3</sup>, 2.960*10<sup>-3</sup>] | 2.552*10<sup>-4</sup> | 9.633 | <.001 |
| SC Distance | -2.039*10<sup>-4</sup> [-2.615*10<sup>-4</sup>, -1.462*10<sup>-4</sup>] | 2.932*10<sup>-5</sup> | -6.954 | <.001 |

## Model 3

Model 3 included independent variables of age, sex, small-worldness, SC total, and mean SC distance, differing from Model 2 by the addition of SC total back into the model in order to investigate the measures of SC total and SC distance concurrently. The results largely mirrored those found in Models 1 and 2. In particular, SC total was positively associated with local efficiency for YA and OA, but this effect was stronger for OA (this group difference was significant based on the 95% CI of the estimate), and SC distance was negatively associated with local efficiency for both YA and OA. Small-worldness was positively associated with local efficiency in both age groups but this relationship was stronger for OA (this group difference was significant based on the 95% CI of the estimate; see Tables 5 and 6).

**Table 5.** Local Efficiency Multilinear Regression Analysis Model 3 for YA, with dependent variable local efficiency mean for regions negatively associated with fluid intelligence based on the PLS analysis. Independent variables included sex, age, small-worldness, SC total (total SC streamline probabilities), and SC distance (mean of all SC streamline distances). Model *R*^2^ = .376.

| Effects | Estimate [95% CI] | Std. Error | *t*-value | *p*-value |
| --- | --- | --- | --- | --- |
| Intercept | .023 [.013, .033] | .005 | 4.501 | <.001 |
| Age | $-3.409 \times 10^{-5}$ [$-5.771 \times 10^{-5}$, $-1.047 \times 10^{-5}$] | $1.199 \times 10^{-5}$ | -2.843 | .005 |
| Sex (Male) | $5.274 \times 10^{-4}$ [$8.237 \times 10^{-5}$, $9.723 \times 10^{-4}$] | $2.259 \times 10^{-4}$ | 2.335 | .020 |
| Small-worldness | $1.464 \times 10^{-3}$ [$1.043 \times 10^{-3}$, $1.885 \times 10^{-3}$] | $2.139 \times 10^{-4}$ | 6.844 | <.001 |
| SC Total | $1.424 \times 10^{-4}$ [$1.106 \times 10^{-4}$, $1.742 \times 10^{-4}$] | $1.613 \times 10^{-5}$ | 8.826 | <.001 |
| SC Distance | $-1.240 \times 10^{-4}$ [$-1.914 \times 10^{-4}$, $-5.656 \times 10^{-5}$] | $3.421 \times 10^{-5}$ | -3.623 | <.001 |

**Table 6.** Local Efficiency Multilinear Regression Analysis Model 3 for OA, with dependent variable local efficiency mean for regions negatively associated with fluid intelligence based on the PLS analysis. Independent variables included sex, age, small-worldness, SC total (total SC streamline probabilities), and SC distance (mean of all SC streamline distances). Model *R*^2^ = .738.

| Effects | Estimate [95% CI] | Std. Error | *t*-value | *p*-value |
| --- | --- | --- | --- | --- |
| Intercept | $-6.018 \times 10^{-3}$ [$-1.487 \times 10^{-2}$, $2.832 \times 10^{-3}$] | $4.500 \times 10^{-3}$ | -1.337 | .182 |
| Age | $6.409 \times 10^{-5}$ [$4.908 \times 10^{-5}$, $7.910 \times 10^{-5}$] | $7.631 \times 10^{-6}$ | 8.398 | <.001 |
| Sex (Male) | $-1.853 \times 10^{-4}$ [$-5.382 \times 10^{-4}$, $1.677 \times 10^{-4}$] | $1.794 \times 10^{-4}$ | -1.032 | .303 |
| Small-worldness | $3.930 \times 10^{-3}$ [$3.491 \times 10^{-3}$, $4.370 \times 10^{-3}$] | $2.234 \times 10^{-4}$ | 17.590 | <.001 |
| SC Total | $1.938 \times 10^{-4}$ [$1.680 \times 10^{-4}$, $2.197 \times 10^{-4}$] | $1.313 \times 10^{-5}$ | 14.763 | <.001 |
| SC Distance | $-1.088 \times 10^{-4}$ [$-1.557 \times 10^{-4}$, $-6.183 \times 10^{-5}$] | $2.386 \times 10^{-5}$ | -4.559 | <.001 |

Taken together with the local efficiency PLS analyses, these results suggest that local efficiency is associated with worse fluid intelligence in OA (as shown by the PLS analysis, see Figure 6), and is also associated with greater connectivity overall in the brain (SC total) and shorter structural connections (mean SC distance; see Table 6). The results also identified a sex effect that differed between YA and OA, whereby males had greater local efficiency than females in YA, but females had greater local efficiency in OA. Greater local efficiency in these regions increases the small-worldness of the brain network, especially for OA, which has been shown to help support organized network function. This suggests that local efficiency may help shore up the structural brain network in response to other factors in the brain that produce declining cognitive ability, and this may be accomplished by short-range connections that support local brain function (in line with Heisz et al., 2015).

## Discussion

We have demonstrated that differences in the whole-brain SC network are related to cognitive ability differences across the lifespan. However, this relationship is nuanced. There are many structural connections throughout the brain that have less streamlines in older adults than in younger adults. This lack of streamlines is associated with declining cognitive abilities, which is expected given the cortical atrophy and demyelination that is known to occur with age. We have also produced the novel demonstration that there are several connections (especially intrahemispheric) that are stronger with age and support spared cognitive function. That these connections are overwhelmingly intrahemispheric supports past research showing that greater functional complexity locally and lower complexity involving long-range interactions (especially interhemispheric) occurs in OA, and that this pattern may be advantageous as it was associated with better cognitive ability (Heisz et al., 2015). Furthermore, we also identified structural connections that were associated with spared cognitive ability in OA when the streamline probability was minimized, suggesting that some of the beneficial configurations of the network involve deprioritizing certain connections in favor of others.

## Nodal Efficiency

By applying graph theory to these networks, we investigated these connectivity changes more comprehensively, using nodal efficiency and local efficiency measures. Nodal efficiency measures how well a region is able to communicate with the rest of the brain and is a marker of hubness. We found that non-typical nodal efficiency is detrimental in many brain regions with typically high nodal efficiency (as shown by mean nodal efficiency in Figure 6), suggesting that there is a relatively ideal network configuration and that alterations to hub-like regions are associated with worse cognitive ability. This seems to be the case primarily for regions including the insula, ventrolateral and dorsolateral prefrontal cortex, medial parietal regions, putamen, cingulate cortex, and caudate nucleus. This suggests that more connectivity is not always better, as this can disrupt the ability for the network as a whole to function effectively. However, especially in OA there are a number of regions, particularly in the RH hippocampus, bilateral nucleus accumbens, and RH ventral occipital-temporal stream, that relate to better cognitive ability with increased hubness as measured by nodal efficiency. The nucleus accumbens has been noted in research on the aging brain to be somewhat unique, in that although volume reductions have been noted with age, it is spared from neuronal loss, unlike much of the rest of the brain (see Konar-Nié et al., 2023 for a review). It may be that greater connectivity bilaterally to this relatively protected subcortical region serves to spare the network, especially considering the role the nucleus accumbens plays in decision making, particularly when there is uncertainty, ambiguity, or when distractors are involved, owing to the role of neurons in this region for integrating information from multiple sources that can be conflicting (see Floresco, 2015 for a review). Frontal and temporal lobe regions provide inputs to the nucleus accumbens, which then prioritizes the most important inputs while dampening less relevant inputs in order to guide responses (Floresco, 2007; Nicola, 2007; O’Donnell, 2003). Greater hubness of the hippocampus was also shown to support cognitive ability. Greater global connectivity of the hippocampus may help to preserve the medial temporal lobe memory system, and counteract its susceptibility to degradation in aging and Alzheimer’s disease (Beason-Held et al., 2021). Furthermore, the hippocampus is involved in spatial processing and relationships between stimuli, and is connected to the nucleus accumbens via excitatory pathways (Floresco et al., 1997; Ito et al., 2008; Mannella et al., 2013). The ventral occipital-temporal stream contains a number of modular regions in the fusiform gyrus that are specialized to some extent for specific visual recognition tasks, such as object recognition (lateral occipital complex; Grill-Spector et al., 2001), face recognition (fusiform face area; Kanwisher & Yovel, 2006), visual word recognition (ventral occipitotemporal cortex; Price, 2012), and place recognition (parahippocampal place area; Aminoff et al., 2013). It may be that the greater hubness and connectivity to the ventral occipital-temporal stream helps to maintain the overall network integration of these regions important for tasks relying on visual processing, which are crucial for success on the fluid intelligence test. Furthermore, the ventral occipital-temporal stream is implicated as far as the anterior temporal lobe, which has been identified as a crucial hub in the semantic memory network that integrates the multiple modalities involved in semantic memory (Patterson et al., 2007). Semantic memory is unique in that it is one of the few cognitive domains that is spared in general as people age, and is supported by functional changes with age (Hoffman & Morcom, 2018). The greater hubness of the anterior temporal lobe with age may be a structural connectivity factor contributing to robust semantic memory in older adults.

These findings can also be contextualized with respect to past research looking at the functional neural correlates of fluid intelligence. Across a range of studies fluid intelligence has been associated with regions in the frontopartietal network (Barbey et al., 2014; Gläscher et al., 2010; Mitchell et al., 2023; Momi et al., 2020; Samu et al., 2017; Smith et al., 2022; Woolgar et al., 2010). We found that YA relied on the mPFC and IPL regions of this network, in line with past research. In comparison the OA adult structural brain network has reprioritized the connectivity of the ventral inferior temporal lobe (also part of the frontoparietal network; Marek & Dosenbach, 2018). Our findings focused on the structural brain network across the lifespan rather than functional activation patterns (as in the research just discussed), and yet we identified similar regions of the frontoparietal network that are implicated with age and fluid intelligence in YA as well as the novel finding that structural connections to the ventral inferior temporal lobe regions of the frontoparietal network take priority in the network at later stages in the lifespan.

The relationship between nodal efficiency and cognitive ability was also investigated across the lifespan using a rolling correlation approach. This analysis revealed an inverted ‘U’ relationship with age, and a critical age window between the ages of 55-72 during which nodal efficiency in regions identified by the OA PLS supports cognitive ability. Strengthened nodal efficiency in these regions may represent a specialized reorganization for supporting cognitive ability that is advantageous for older adults, owing to the unique architecture of the healthy aging brain network. Ultimately though, the benefits of this reorganization drop-off over time as the critical window is passed, highlighting the importance of interventions that target this subnetwork early enough in the lifespan.

## Local Efficiency

Local efficiency, an indicator of fault tolerance that measures how well connected a region’s neighbors would be if it was removed from the network, was found to be positively associated with cognitive ability in YA for many regions in the brain including cortical regions such as extrastriate cortex, precuneus, retrosplenial cortex, parrahippocampal cortex, medial parietal cortex, temporal and temporoparietal cortex, insula, auditory cortex, and precentral and postcentral gyri, as well as the hippocampus and thalamus in the subcortex. However, the pattern is completely reversed for OA, in which brain regions show a negative association between local efficiency and cognitive ability. By investigating the effect that local efficiency differences have on the whole-brain connectivity patterns, we found that the local efficiency of these regions is associated with greater small-worldness of the brain network throughout the lifespan, but especially so in OA. In OA, greater local efficiency may occur in response to factors related to declining cognitive ability, and may use short-range connections to achieve this network configuration. As with nodal efficiency, the rolling correlation analysis of local efficiency and cognitive ability revealed an inverted ‘U’ relationship with age and a critical window. However, in this case the critical window was earlier in the lifespan for middle-aged adults between 32-51 years old, suggesting that the ideal window for interventions may be even earlier than indicated by nodal efficiency.

## Limitations and Future Directions

It is important to note that this work is based on the cross-sectional data available to us, rather than the more ideal use of longitudinal data when available. Indeed, recent research has highlighted that there are important differences in what is being measured with cross-sectional and longitudinal data (Vidal-Pineiro et al., 2021). However, we have benefitted greatly from the quality of the Cam-CAN dataset and the ability to observe the lifespan changes in how SC relates to age and cognitive ability over such a large range of ages, from the beginning of adulthood (age 18) to late in the aging process (age 88). Future research with longitudinal data covering a similar age range should investigate these effects from a longitudinal perspective. Furthermore, an investigation of how the SC network grows to support cognitive ability in children and adolescents would contribute to a fuller understanding of how the plastic SC network evolves throughout the entire lifespan.

## Conclusion

Whether changes in the functional strategies used by the brain are driving the structural connectivity changes we observed because of the increased use of certain structural connections, or whether the reverse is true, whereby structural connectivity changes are produced in order to encourage new functional strategies, we have demonstrated that healthy aging is not simply a matter of functional adaptations to structural deficiencies. Our findings suggest that not all structural changes are detrimental with age, but rather that functional *and* structural changes may occur to spare cognitive ability in older adults. Rather than needing to keep the brain youth-like to maintain cognitive ability, there are multiple ways in which the healthy aging brain represents a unique structural network regime that is distinct from what is ideal for younger adults.

## Acknowledgements

This research was supported by the Natural Sciences and Engineering Research Council of Canada (NSERC) through Postdoctoral Fellowships Program funding to Josh Neudorf, and by NSERC Discovery Grant *RGPIN-2018-04457* and Canadian Institutes of Health Research (CIHR) Project Grant *PJT-168980* to the senior author Anthony R. McIntosh. This research was enabled in part by support provided by the British Columbia DRI Group and the Digital Research Alliance of Canada (alliancecan.ca). The authors affirm that there are no conflicts of interest to disclose.

## Supplementary Materials

**Supplementary Figure 1.**
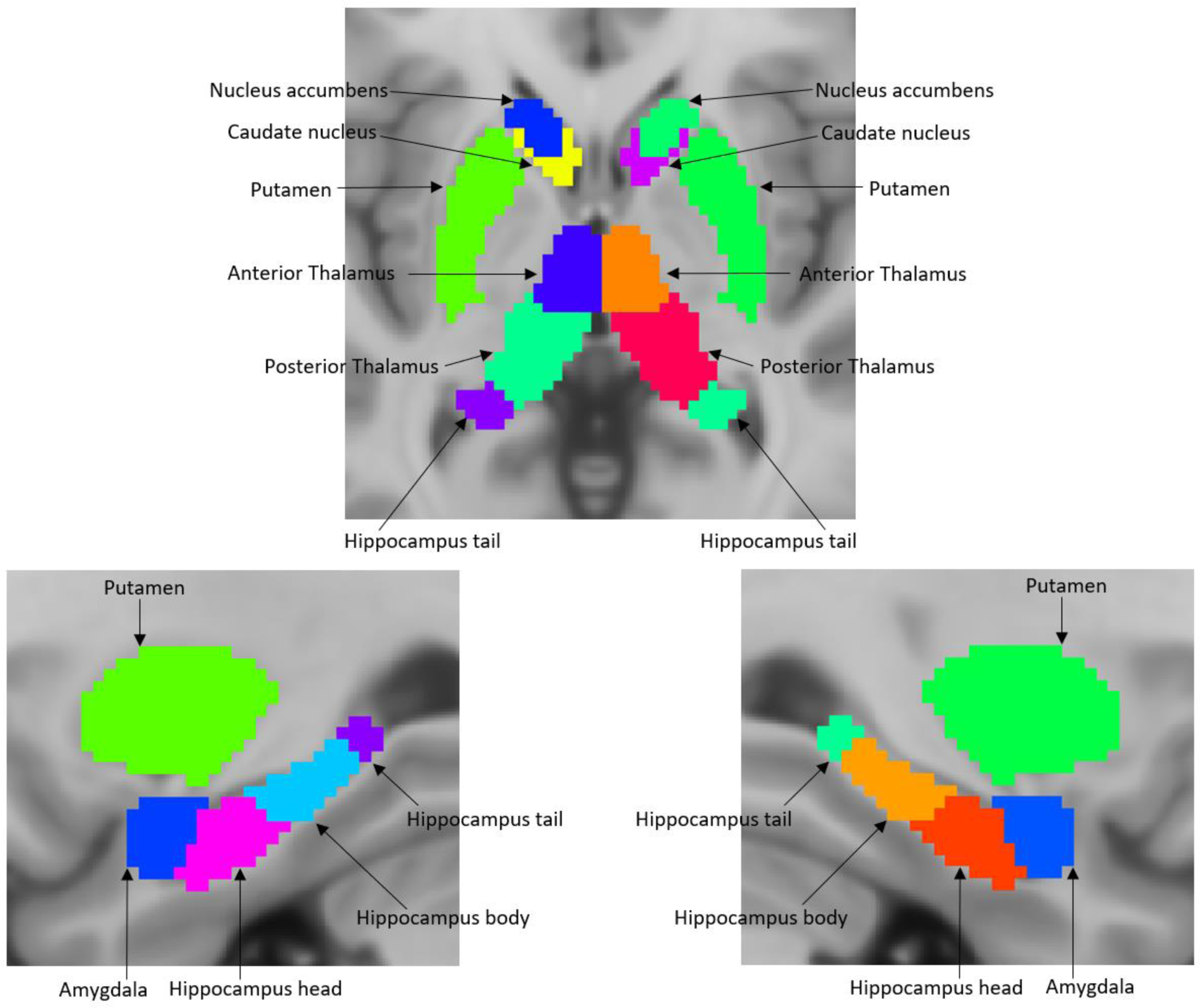
Tian et al. (2020) subcortical parcellation labels.

**Supplementary Figure 2.**
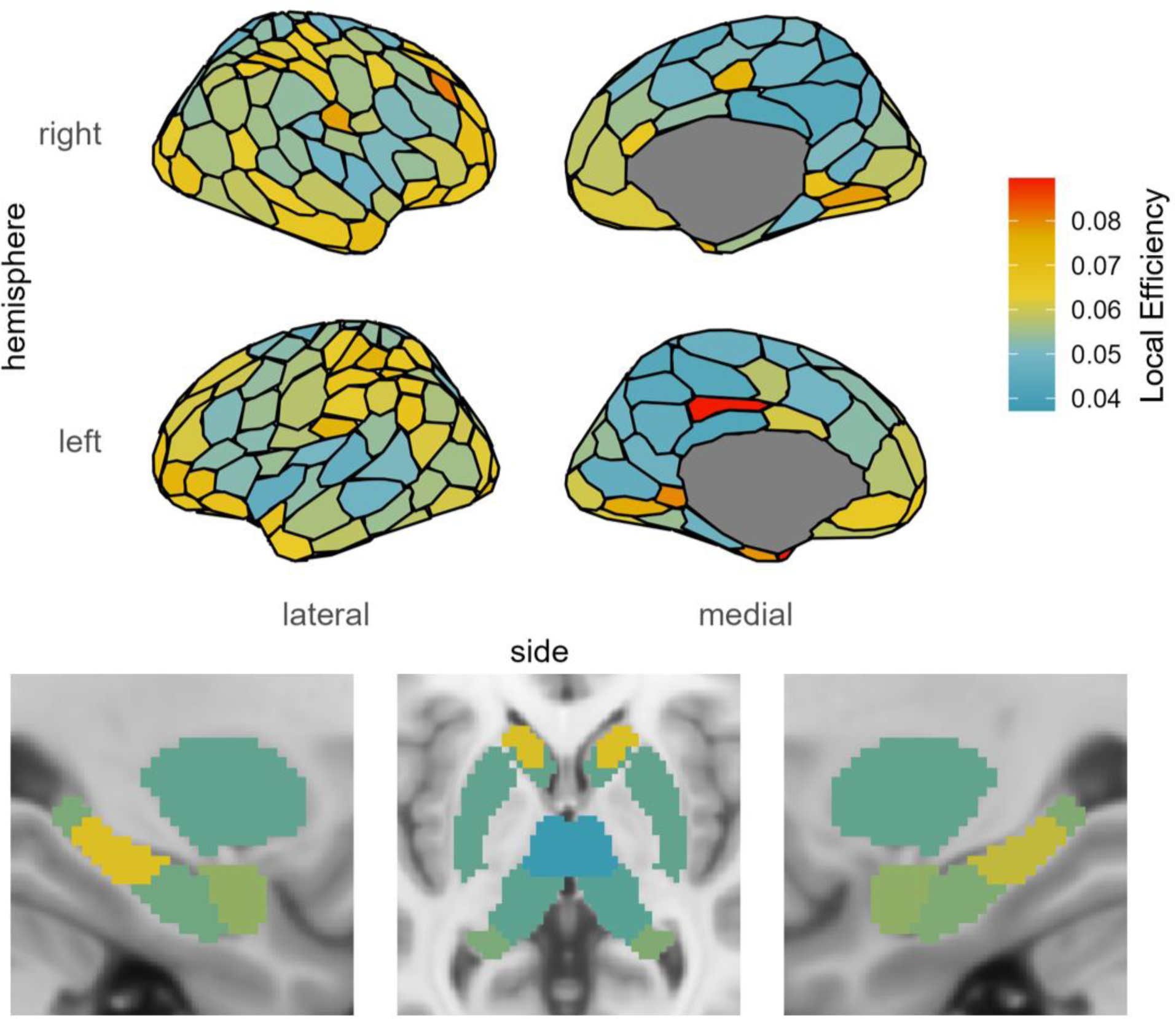
Brain map of mean local efficiency for all participants.

